# AQMM: Enabling Absolute Quantification of Metagenome and Metatranscriptome

**DOI:** 10.1101/218347

**Authors:** Xiao-Tao Jiang, Ke Yu, Li-Guan Li, Xiao-Le Yin, An-Dong Li, Tong Zhang

## Abstract

Metatranscriptome has become increasingly important along with the application of next generation sequencing in the studies of microbial functional gene activity in environmental samples. However, the quantification of target active gene is hindered by the current relative quantification methods, especially when tracking the sharp environmental change. Great needs are here for an easy-to-perform method to obtain the absolute quantification. By borrowing information from the parallel metagenome, an absolute quantification method for both metagenomic and metatranscriptomic data to per gene/cell/volume/gram level was developed. The effectiveness of AQMM was validated by simulated experiments and was demonstrated with a real experimental design of comparing activated sludge with and without foaming. Our method provides a novel bioinformatic approach to fast and accurately conduct absolute quantification of metagenome and metatranscriptome in environmental samples. The AQMM can be accessed from https://github.com/biofuture/aqmm.

## Background

Shotgun metatranscriptomics is a powerful tool in identifying the overall expression of microorganisms in an environment (Alexander et al. 2015, Gifford et al. 2011, Shi et al. 2009, Turner et al. 2013, Yu and Zhang 2012), shedding light on discovering how microbes respond to environmental changes or diseases status (Jorth et al. 2014, Mason et al. 2012) and capturing gene expression patterns for functionally important bacteria in engineering systems (Oyserman et al. 2015, Stark et al. 2014). For these applications, accurate quantification is required to detect the true variations or differential expression genes (DEGs).

Traditionally, the abundance of a transcript in RNA-sequencing (RNA-seq) is thought to be influenced by the library size and inherent dependence on the expression levels of other transcripts as described in a comprehensive review (Rapaport et al. 2013). Following this idea, transcripts in RNA-seq was generally quantified by within-sample normalization. One of the most common quantification methods was RPKM (Mortazavi et al. 2008) (reads per kilobase of exon model per million mapped reads) which considered factors of both the length of gene and library size. Another improved within-sample normalization method was TPM (transcript per million) (Wagner et al. 2012) which only considered the transcript rather than the whole library size and respected the invariance of relative molar RNA concentration (rmc). The TPM was thought to be better fitted in sample comparison due to its unit-free characteristics. The FPKM (substitute the reads with fragments in RPKM) was an adaption of RPKM to pair-end reads. These above methods are all relative quantification (RQ) and suffer from the ‘composition effects’ (the increase of one transcript will decrease other unrelated transcript). To relieve this problem, Robinson and Oshlack proposed a new normalization method “TMM” (trimmed mean of M-values) to detect the DEGs under the hypothesis that most of the genes are not differentially expressed (Robinson and Oshlack 2010), which has been integrated into popular DEGs detection R software edgeR (Robinson et al. 2010). The scaling factor in edgeR for normalization is the TMM value. Another method was to compute the median of the ratio as the scaling factor and it could be conducted by R software DESeq/DESeq2 (Love et al. 2014). It is also based on the assumption that most genes are not DEGs and this method then calculates the scaling factor (median of ratios) associated with this sample to perform further normalization. In the two software, the negative binomial distribution was applied to adjust the distribution of transcript between different conditions to relieve the dispersion effects of deviation from standard passion distribution (Rapaport et al. 2013). Although with these efforts in optimizing the normalization process, these indices were all still RQ based and the relationship could be distorted while performing comparative analysis across samples, especially when borrowing these methods from traditional Eukaryote RNA-seq to current Prokaryote metatranscriptome studies (Conesa et al. 2016). One feasible way to solve the problem was to get the absolute quantification (AQ) of expression level for each transcript. For example, the qRT-PCR has long been applied in RNA-seq or microarray data for AQ (Becker-André and Hahlbrock 1989, Whelan et al. 2003). In addition, there were methods by spiking in exterior/alien RNA in microarray to get the per cell absolute quantification (Kanno et al. 2006) and internal standard approach to estimate per liter expression in marine metatranscriptome (Gifford et al. 2011). However, the experiment to perform spiking internal standard was difficult due to its skill-demanding nature and for metatranscriptome data, factors like the time to add spike-in material, the type and the amount of alien RNA required still needed to be elaborately designed. Hence, it was not as popular as those RQ methods. The quantification methods in the newly developed analyzing pipelines for metatranscriptome like IMP (Narayanasamy et al. 2016), MetaTrans (Martinez et al. 2016), COMAN (Ni et al. 2016) and SAMSA (Westreich et al. 2016) were still all based on RQ methods; this would result in accelerated spreading of the inaccurate quantification in many studies.

To solve the problem of RQ and get an accurate quantification without performing spike-in experiment, an AQ bioinformatics software package AQMM was developed by combining metagenome and metatranscriptome data to achieve the goal of accurate and comparable quantification. In this study, we firstly introduced the AQMM algorithm flow, and then compared and validated it with RQ methods by simulated metagenome and metatranscriptome data. Moreover, we further applied this algorithm to a real combination of metagenome and metatranscriptome dataset in quantifying genes and transcripts of resistome in six foaming activated sludge (FAS) and non-foaming activated sludge (NFAS) samples.

## Results

### Overall view of AQMM algorithm

The AQMM **(Fig. 1)** was designed to perform AQ of parallel metagenome and metatranscriptome dataset no matter whether spike-in experiment/internal standard was initially added or not. The major aims were to obtain the AQ of genes/transcripts/taxa in samples and to accurately detect DEGs in metatranscriptome data. The assumptions under the algorithm include: 1) with the known extraction ratio of DNA for a DNA extraction Kit for a type of sample, the total weight of DNA per volume of the sample could be calculated. The weight of the sequenced library of DNA could be estimated with the molecular weight and bases numbers of A, T, C and G in the sample. Then, the ratio of sequenced DNA to total weight of DNA per volume of the sample could be calculated. In addition, by utilizing the universal single-copy phylogenetic marker genes (USCMGs), the number of cells for a metagenome library could be estimated accurately (Nayfach and Pollard 2015). With the above information, cells per volume could be calculated for a metagenome data; 2) Using the same volume of the sample contains the same number of cells for DNA and RNA extraction, the cell number per volume to extract RNA was the same as the parallel DNA sample; 3) With the known ratio of RNA extraction and the rRNA ratio of total RNA, by a similar process, the sequenced RNA weight ratio could be calculated, and then the equivalent cell numbers in a metatranscriptome could be deduced accordingly. With the cell numbers included in the metagenome and metatranscriptome data, the abundance of gene/transcript could be normalized to per cell level. Moreover, as the number of cells per volume is available, per cell quantification could be easily transformed into per volume quantification.

**Fig. 1:**
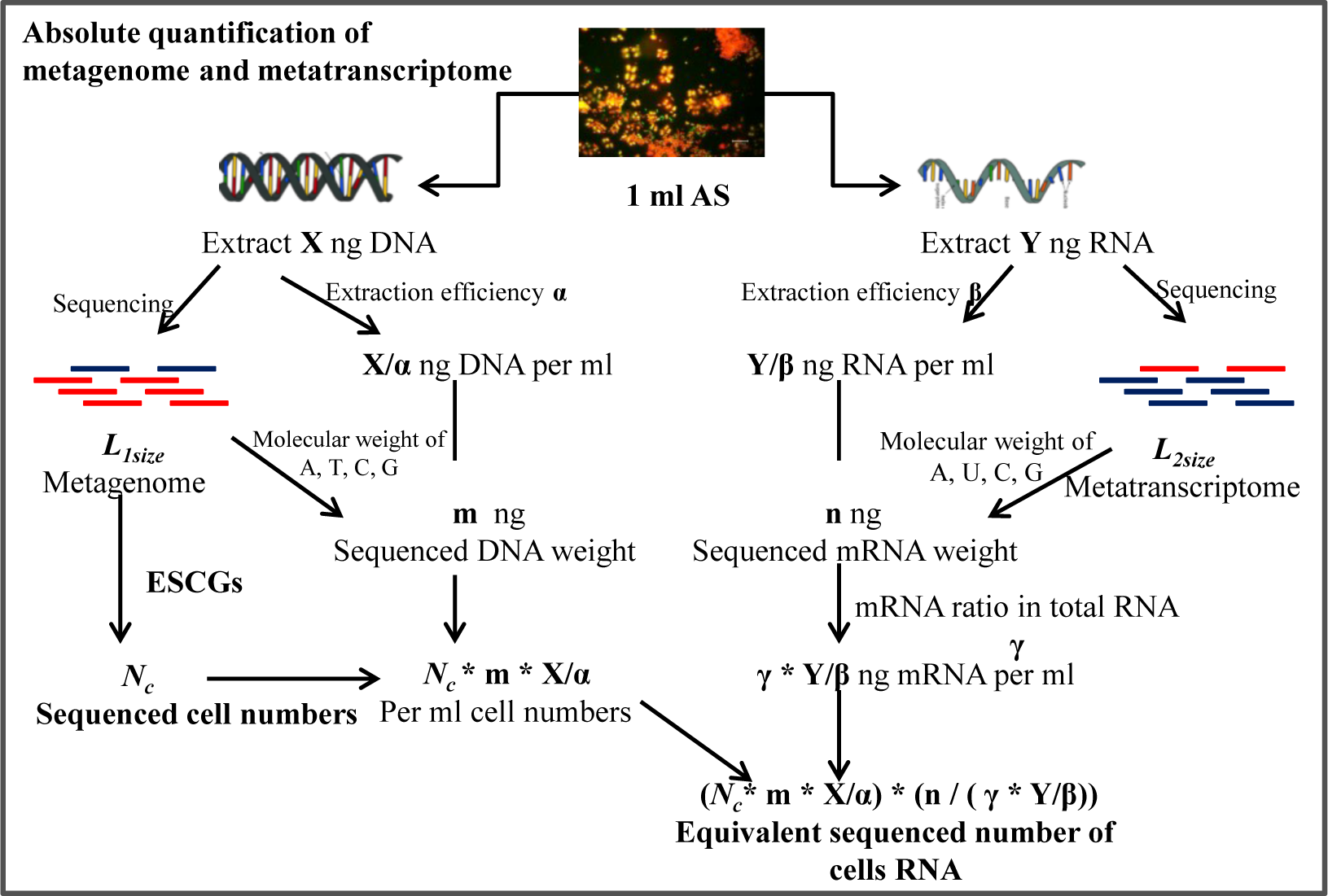
Schematic flow diagram for absolute quantification of metagenome and metatranscriptome to cell/volume level.

### Comparing and validating AQMM with RQ methods using simulated metatranscriptome demo

To reveal the problem of RQ methods like RPKM, edgeR and DESeq2 and to assess the effectiveness of AQMM, simulated metatranscriptomic datasets comprised of known community structure and expression levels were generated **(Fig. 2; Details in methods).** The simulated data was with known ground truth absolute expression for each gene. For simplicity, to focus on the quantification of metatranscriptome in identifying DEGs, we assume the DNA content are not changed like what happens in a reactor with a stable biomass concentration, however the gene expression under condition A and B are significantly changed with fold of 2 or 16 in part of the bacteria like what happens under sharp environmental change. In order to focus only on the of influence normalization methods, in generation of the simulated metatranscriptome, the base qualities of were all set with 50 and to eliminate the influence of mapping process, the mapping criteria of bowtie2 was set to exactly match without gap and mismatch allowed (bowtie2 parameters, ‐N 0 ‐L 31, ‐‐rdg 100,150 ‐‐rfg 100,150 ‐‐gbar 100,150). The result of DEGs detection was in **Table 1**. We can observe that compared with ground truth, the RQ methods detect quite a large portion of false positive higher gene expression under condition A. On the contrary, the AQMM method which aims to obtain the AQ has limited errors detection even with a given variance in RNA extraction efficiency **(Table 1)**. Noticeably, in real combination of metagenome and metatranscriptome, the metagenome could also be totally different, and in this case, the AQMM is still applicable.

**Fig. 2:**
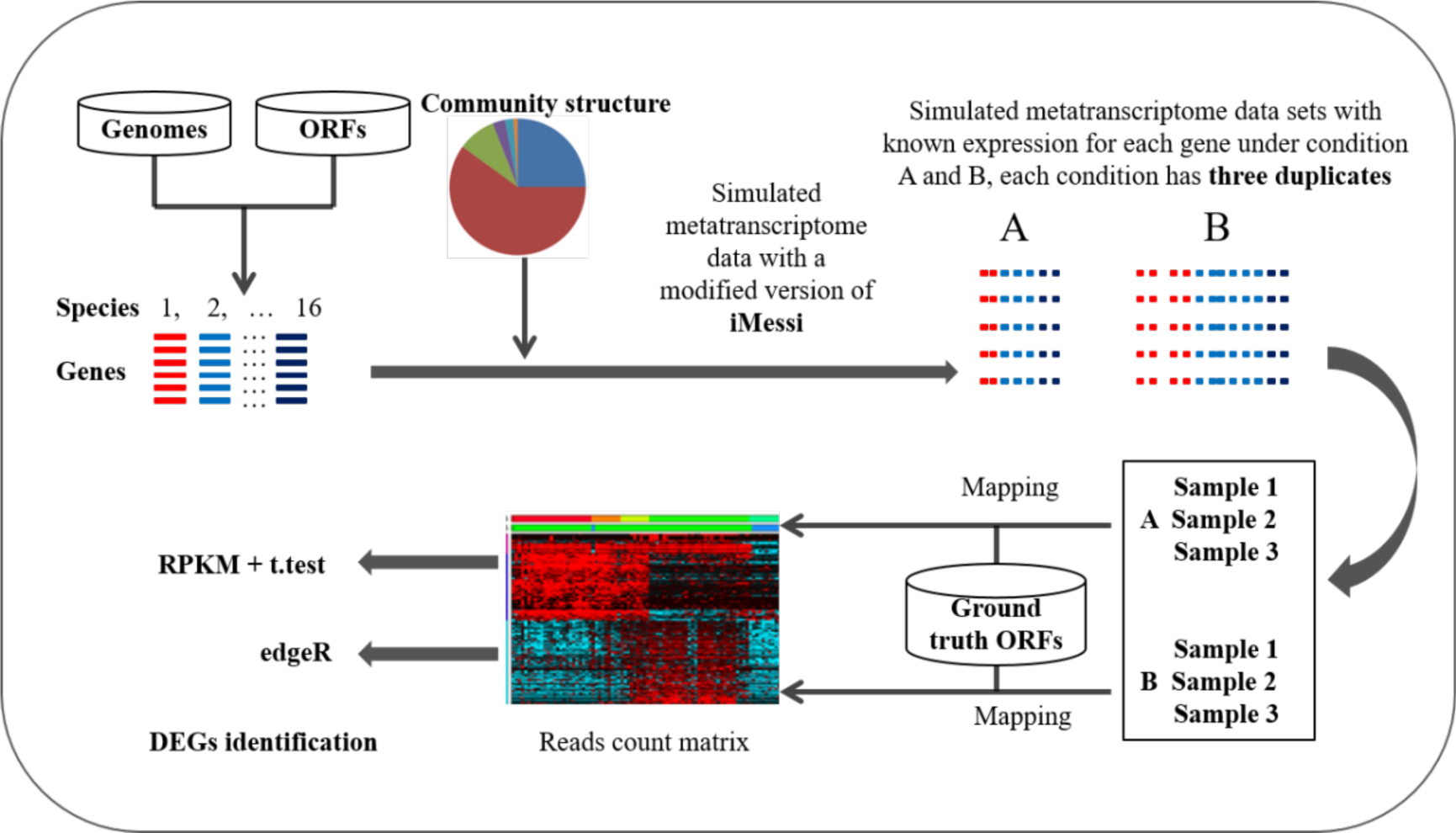
Flowchart of the simulation datasets generation and analyzing process to get the differential expression genes.

**Table 1:** Comparing relative quantification methods with AQMM on detection of DEGs for simulated metatranscriptome data.

|  | # of genes Higher<br>expression in B | No expression<br>difference | # of genes<br>Higher<br>expression in A |
| --- | --- | --- | --- |
| **Theoretical Ground Truth** | **28524** | **36572** | **0** |
| **RPKM+t-Test (P < 0.05)** | 16477 | 11558 | **37062** |
| **edgeR** | 18278 | 20778 | **26040** |
| **AQMM-5%-variation** | 28744.72 ± 143.53 | 35807.52 ± 48.08 | **543.77 ± 129.72** |
| **AQMM-10%-variation** | 28740.83 ± 298.43 | 35801 ± 188.31 | **554.17 ± 256.81** |
| **AQMM-20%-variation** | 28549.48 ± 1007.17 | 35941.86 ± 919.86 | **604.66 ± 654.76** |
| **AQMM-50%-variation** | 16673.93 ± 9394.27 | 47694.99 ± 9600.33 | 727.08 ± 1775.09 |

### Case study: AQ of activate resistome in FAS and NFAS

The AQMM was applied in the six metagenome and metatranscriptome dataset of FAS and NFAS, the AQ of the sequenced cells generated by the pipeline were shown in **Table 2**. In detail, the metagenome contained 8 to 11.8 GBs data and metatranscriptome with a depth between 13 and 16 GBs for each sample. The “per cell/volume” quantifying values were the fundamental of normalizing to cells or volume in order to perform comparison among different studies. The cell number per milliliter in literature was at 3.3E+09 using flow Cytometer to quantify(Foladori et al. 2010) and was from 2.1E+09 to 5.5E+09 using CFU and flow Cytometer (Manti et al. 2008) level for AS which was a bit lower than the obtained number in this study at the magnitude of E+10 cells per milliliter. Overall number of mRNA molecules per cell are 387.98 ± 102.86 and 235.21 ± 30.59 averagely for FAS and NFAS, respectively **(Table S1)**, which is consistent with previous observation of coastal bacterioplankton by 142-238 mRNA molecules per Cell (Gifford et al. 2011, Moran et al. 2013).

**Table 2:**
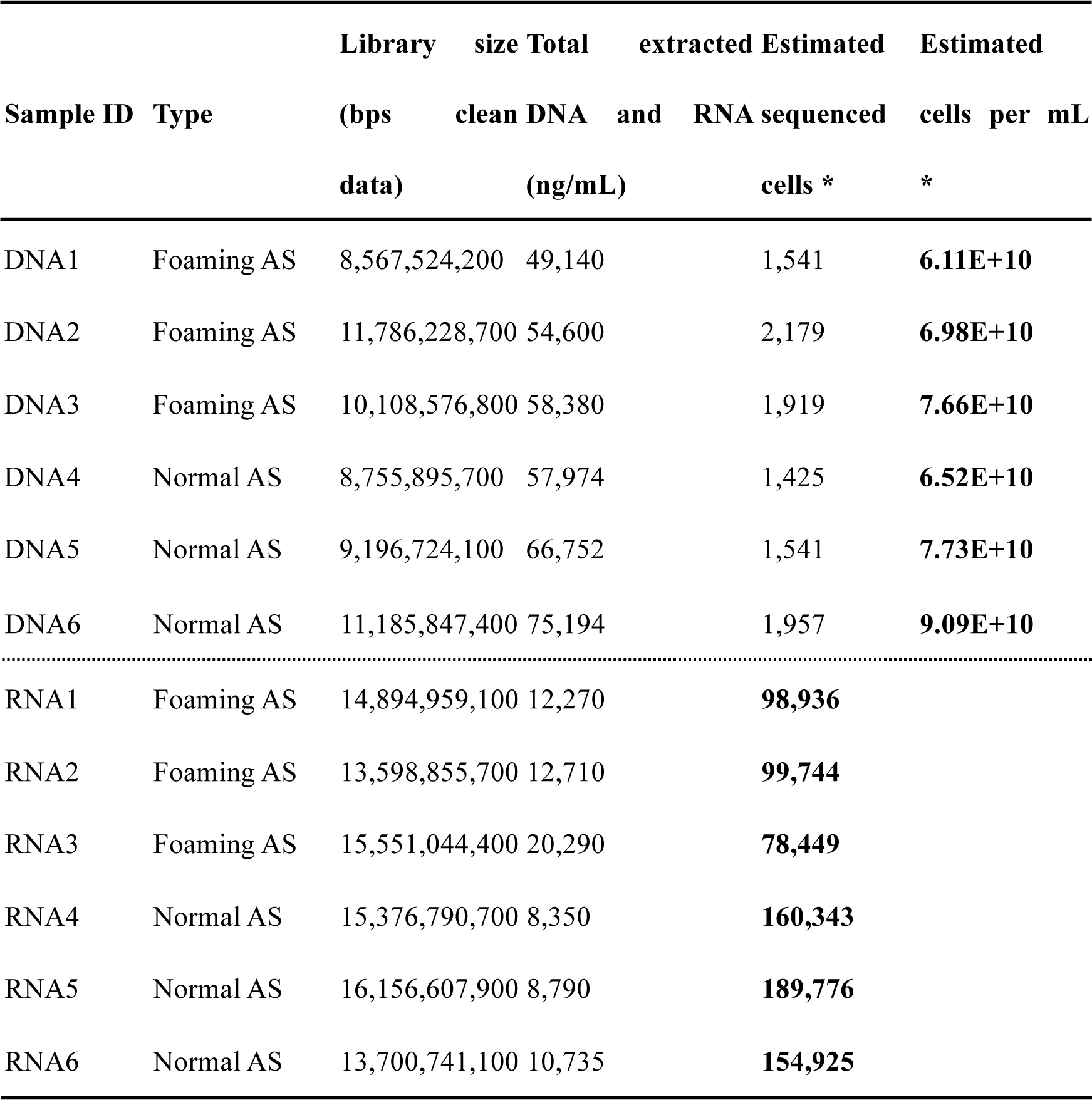
Summary of sequencing outputs and absolute quantification of each sample at cell level with AQMM.

As WWTPs become the hot-spot of antibiotic resistant genes (ARGs) to the receiving environment. Hence, the expressions of ARGs in the AS were in great concerns and further profiled. Overall, the abundance of ARGs per cell in FAS and NFAS were 0.0517 ± 0.0034 and 0.0483 ± 0.0041; and the transcript of ARGs per cell were 0.0140 ± 0.0039 and 0.0059 ± 0.0009, respectively **(Table S2 & S3)**. The overall transcription of ARGs was significantly higher in FAS compared with NFAS. At DNA level, only tetracycline resistance gene was higher in FAS and beta-lactam was higher in NFAS, other types were not significantly different. However, at transcript level, all the types were all significantly higher in FAS. Among the nine transcribed ARGs types, beta-lactam and sulfonamide resistance genes were the most abundant expressed ARG types in both FAS and NFAS. Per volume ARGs abundance and expression at type level were shown in **Fig. 3**. The overall ARGs abundance per milliliter AS in FAS and NFAS were 2.51E+09 ± 2.44E+08 and 2.66E+09 ± 5.63E+08; and the transcript of ARGs per milliliter were 9.83E+09 ± 3.82E+08 and 4.49E+09 ± 5.10E+08, respectively. With the AQ results, the transcripts per copy gene (TPCG), which represents of the transcribe rate could be further derived. The unclassified, quinolone, multidrug and beta-lactam were more active in FAS compared with NFAS in terms of TPCG, **(Table S4)**. For the detected ARGs, the host taxonomy was assigned by LCA algorithms using all the genes annotation in the same Contig. Thirteen orders were detected to carry ARGs and eleven of them were transcribed (**Fig. 3)**. The most ARGs transcribed order was Enterobacteriales. The active ARGs in bacteria enclosed in foams of FAS posed potential threats for the public as ARGs carrying bacteria could spread into the air from the foams bubbles.

**Fig. 3:**
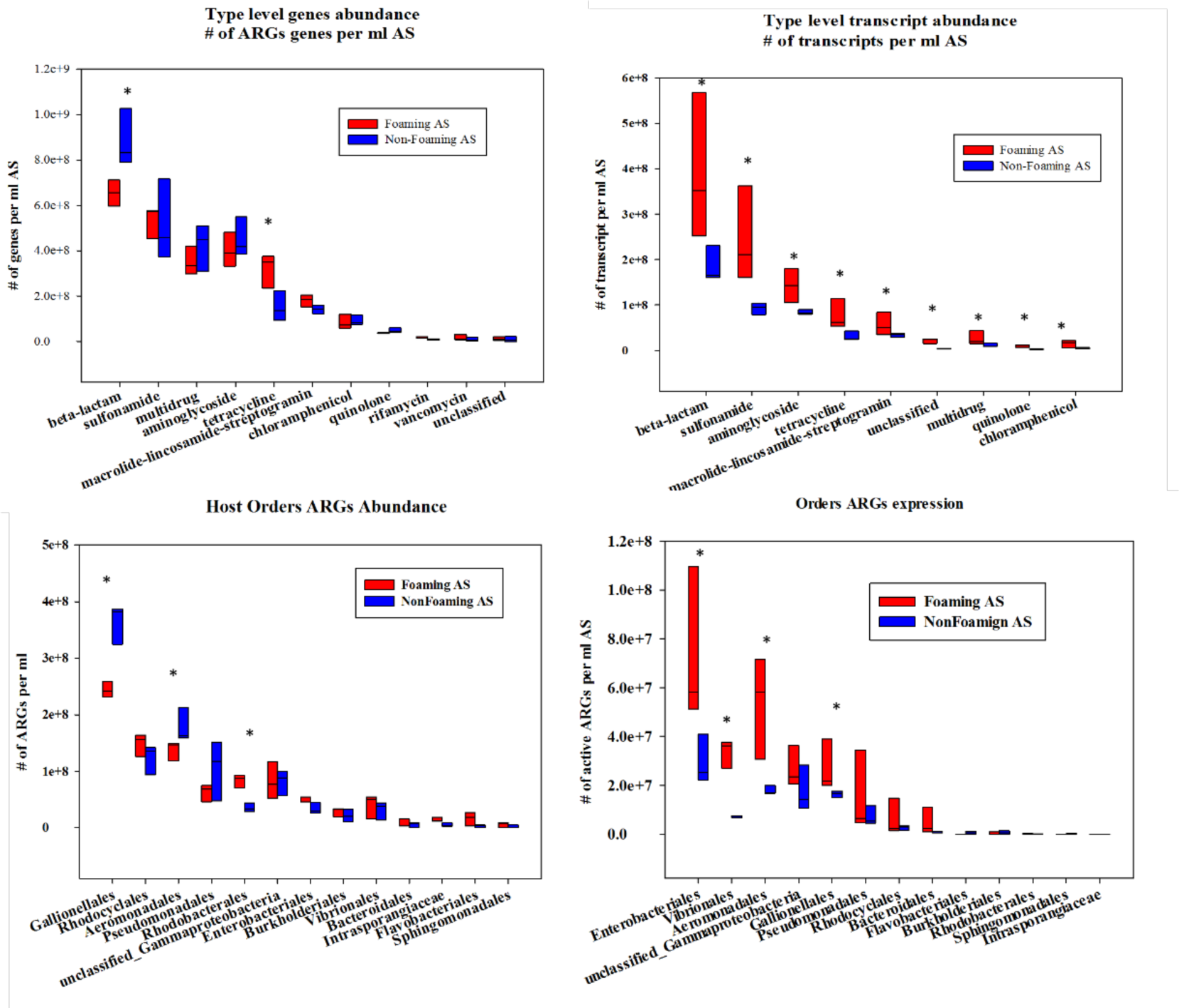
Absolute quantification of type level ARGs abundance and transcription in FAS and NFAS. ARGs-carry hosts abundance and expression. * represents significant difference (*P*-value < 0.05).

The co-expression of ARGs and MRGs was also studied to check whether there were co-expression effects at the RNA level. Using this dataset, we observed co-expression within ARGs, within MRGs, and between ARGs and MRGs (**Fig. 4**). Numerous types of MRGs were detected in the metagenome and metatranscriptome. The most abundant MRG was Cu resistant genes and for the ARGs, beta-lactam, tetracycline and aminoglycoside were the most expressed types. The highest number of co-expression within MRGs was Cr and Fe; while within ARGs was beta-lactam and tetracycline. The most MRG and ARGs co-expression was Cr, which co-expression with nine types of ARGs. This was the first transcript level evidence of the co-expression of ARGs and MRGs in AS.

**Fig. 4:**
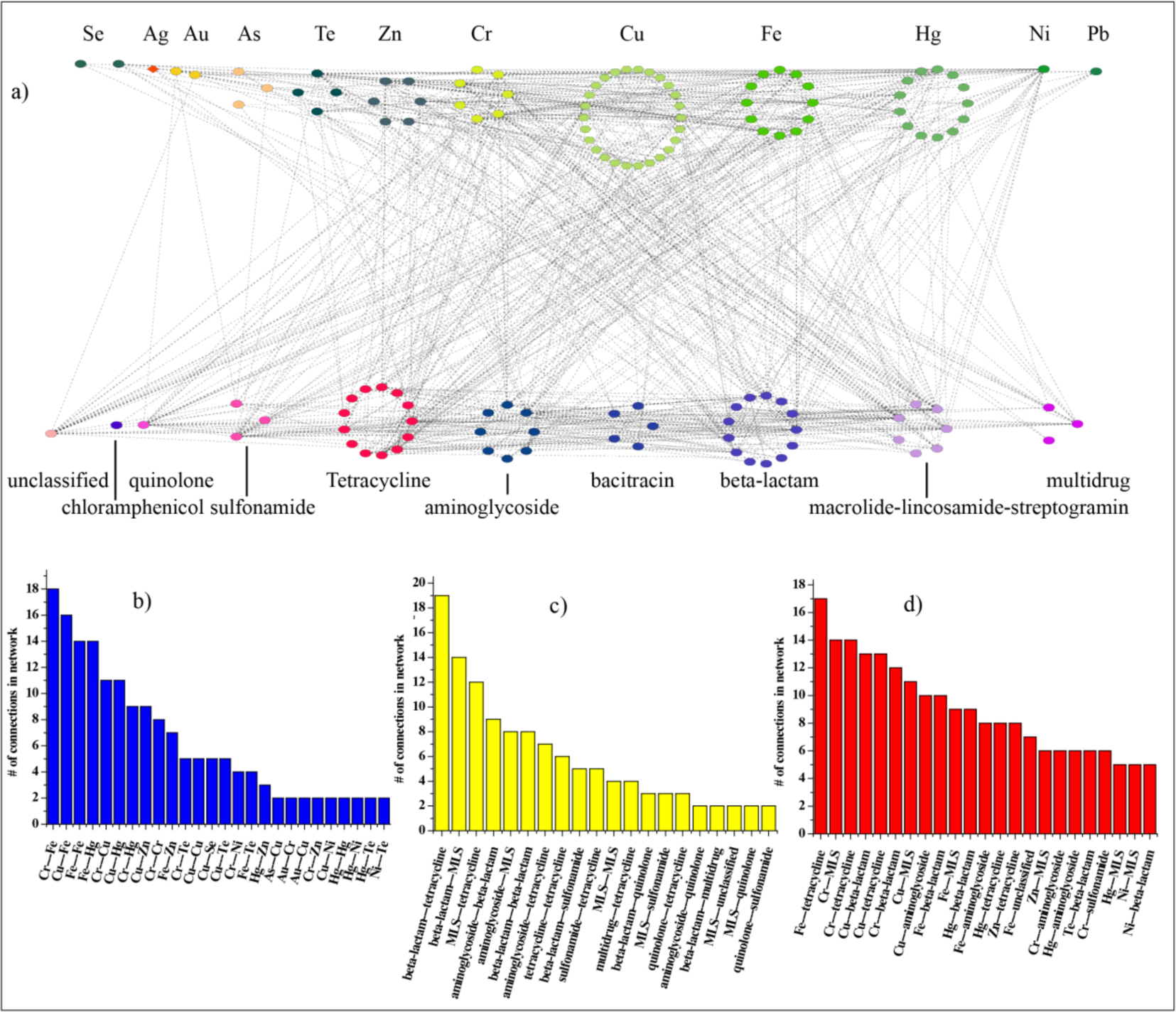
Co-expression of ARGs and MRGs in Shatin AS, a) was the network of ARGs and MRGs expression; b) was statistical of co-expression within MRGs; c) was statistical of co-expression with ARGs; d) was statistical of co-expression of ARGs and MRGs. Lines in the network represented Spearman association over 0.6, *P-*value 0.05 the *P-*value was adjusted with B-H method.

## Discussion

Metatranscriptome enabled the study of whole metabolic pathways expression of the system and many studies had already taken this advantage for different environments, such as in marine (Mason et al. 2012), rhizosphere of the plant (Turner et al. 2013), human oral disease (Jorth et al. 2014). Each study has specific method to integrate the metagenome and metatranscriptome information to understand the microbes and their activities in the system. The quantification of metatranscriptome was generally RQ based methods. The RQ methods are problematic as they may not be able to reflect the actual expression level of a population in the whole community. Due to the relative characteristics, the RQ methods are always suffer from the so-called composition effects, which indicates that the upgrade of one gene should definitely make other genes downgrade. Additionally, the RQ methods are just a relative portion rather than a value with biological implications. On the contrary, the AQ could be more biological meaningful at per cell/volume unit. Hence, it was necessary to conduct AQ to compare different samples. In this study, we proposed an AQ method and developed a set of algorithms to conveniently calculate the absolute number of sequenced cells for each RNA library by borrowing cell numbers from a corresponding data set of DNA library of the same sample.

Noticeably, there were several hypotheses for the application of the proposed method. Firstly, the sample used to extract DNA and RNA should contain the same cell numbers per volume which could be easily met with sufficient mixing of samples. Secondly, the DNA and RNA extraction efficiency should be estimated, as well as the rRNA ratio in total RNA. This was likely difficult to achieve. However, for an environmental sample, generally literature based data could be used for the extraction kit, for example, to FastDNA SPIN Kit for Soil, the extract efficiency was estimated as 28.4% (Mumy and Findlay 2004). Most importantly, as the parallel samples were extracted under the same condition, the difference between samples was minimized **(DNA extraction data, unpublished)**. This AQMM method is capable of performing absolute quantification of both metagenome and metatranscriptome without the requirement to do complex spike-in experiments. Importantly, AQMM avoids the RQ problems of composition effects and able to detect accurate DEGs. Hence, the proposed AQMM is a method in between experimental spike-in based AQ methods and those improved RQ methods of TMM based edgeR.

With AQ, a number of indices with various biological meaning were proposed in this study (Methods), for example, the transcript per copy gene (TPCG) index is a reflection of the transcribe rate of the gene, which could never be delivered by RQ methods. It was demonstrated with simulating RNA-seq that the organism abundance (community structure) was important at normalizing metatranscritptome data in identifying DEGs (Klingenberg and Meinicke 2017). The gene per cell (GPC) and transcript per cell (TPC) in AQMM are global level normalization indices and the scaling factor is the total number of cells in the DNA or RNA library. This global scaling factor could be easily transformed into taxa specific scaling factors with the relative quantification of different taxa with indices of transcript of taxon A per cell (TTPC). Hence, the normalization in AQMM is well fit for the factor of microbial abundance in metatranscriptome data.

AS is important biological wastewater treatment process and this system is considered as a hot spot for ARG dissemination into the receiving water. The foaming of AS would result in spreading of foams with AS bacteria into the surrounding environment. Understanding the active resistome and the host bacteria in foaming AS enables engineers understanding the risk of sludge foaming incurred to the surrounding environment. We observed a wide profile of active ARG types in the FAS, the identification of opportunity pathogen bacteria *Pseudomonas* carrying active ARGs alerts us the risk of spreading ARGs-carrying bacteria. Additionally, per cell mRNA molecules is an important indication of the activity of the cell, generally natural bacterial communities was observed to hold a lower inventory of transcripts (Moran et al. 2013); and the absolute quantification obtained with AQMM was well-fitted with previous observation.

## Conclusions

In this study, we filled the gap of lacking a bioinformatic algorithm to perform AQ of metatranscriptomic data. The developed AQMM was demonstrated to gain enhanced performance at identifying DEGs compared with those RQ methods benchmarked with simulated metagenomic and meatranscriptomic data. Additionally, with the AQMM, the active resistome in foaming and normal activated sludge were quantified to per cell/volume level and even down to the transcription per copy gene. The active ARG host were quantified and the co-expression of MRGs and ARGs was revealed for the first time in AS.

## Materials and methods

### Absolute quantification of gene abundance and transcript expression

We developed a package of scripts AQMM (absolute quantification of metagenome /metatranscriptome) to perform comparative analysis.

The formula for cells per mL:

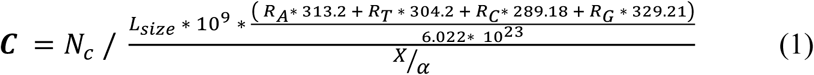

*C* is value of cell numbers per mL AS

*N_c_* is the estimated cell numbers for the sequenced DNA library with USCMGs

*L_size_* is the sequencing depth

*R_A_*, *R_T_*, *R_C_* and *R_G_* are ratios of A, T, C and G

X is the overall extracted weight (ng) of DNA for 1 mL AS

α is DNA extraction efficiency, for FAST DNA Kit for Soil, α is estimated as 28.2% (Mumy and Findlay 2004).

The sequenced cells for RNA sequencing, for a RNA-seq with library size of *L_size_* after removing all ribosomal RNA, the equivalent sequenced cells for this sample is

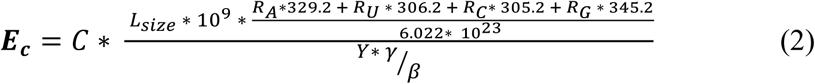

*E_c_* is the estimated number of cells sequenced for this RNA library *C* is value of cell numbers per mL AS *L_size_* is the sequencing depth *R_A_*, *R_U_*, *R_C_* and *R_G_* are ratios of A, U, C and G, the value they multiplied are molecular weight Y is the overall extracted weight (ng) of RNA for 1 mL AS β is RNA extraction efficiency, the estimated β is about 7.5% as used in this study.

This value was deduced from AS empirical data of proportion of RNA biomass by engineering perspective and the extracted RNA biomass.

γ is non-ribosomal RNA ratio, for AS the estimated γ is about 0.03.

Based on the two AQ numbers of cells for each sample, the gene or transcript abundance matrix could be further normalized into the following indices.

**GPC** (Gene per Cell): an indication of the overall abundance of the gene in system.

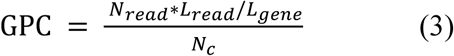

**TPC** (Transcript per Cell): an indication of overall activity of the gene in system.

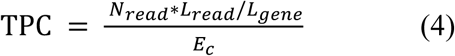

**TPCG** (Transcript per copy gene): an indication of the absolute activity of one copy gene in the system, equivalent to transcribe rate for each gene.

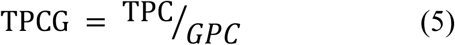

**GTPC** (Gene of taxon A per Cell): an indication of the overall abundance of the taxon in system averagely.

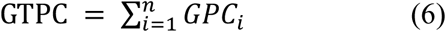

**TTPC** (Transcript of taxon A per Cell): an indication of overall activity of the taxon in system averagely.

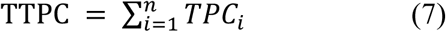

**ATCT** (Averagely transcript per copy gene of taxon A): indication of the averagely absolute activity per copy expressed gene in taxon A

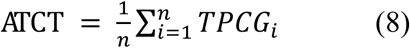

*N_c_* is the estimated cell numbers for the sequenced DNA library, *N_read_* is the number of reads or transcript mapping to the target gene *L_read_* is the length of reads *L_gene_* is the length of the target gene n is the number of genes affiliated to taxa A.

When the number of cells per mL was obtained, using the GPC, genes per mL could be calculated.

### Simulating metatranscriptome data

To validate our method and comparing with those RQ methods in identifying the DEGs, simulated data was generated by workflow illustrated in **Fig. 2**. For simplicity, the DNA was set unchanged to mimic the activated sludge community composition with 16 strains from different phylogeny. The metatranscriptome data sets were generated for two conditions A and B, each with three biological duplications; for the condition A and B, there were part of the strains with folds of significantly changed expression (**Table S5**). To only focus on the quantification method, all the system errors caused by other factors like base qualities, cDNA synthesis, assembly, mapping parameters were not considered.

### Sampling

AS samples were collected in Shatin wastewater treatment plant at three locations along the flow direction while serious foaming happened at 2016-04-08 and nearly no foaming happened at 2016-04-25. Samples were collected on site by storing in liquid nitrogen immediately and then transported to the laboratory for RNA extraction. The DNA samples were mixed with 1:1 100% ethanol and AS and then stored at −20 °C fridge. Totally six samples were collected for both DNA and RNA samples alongside the segment aeration tank in three locations as depicted in **Fig. 5**.

**Fig. 5:**
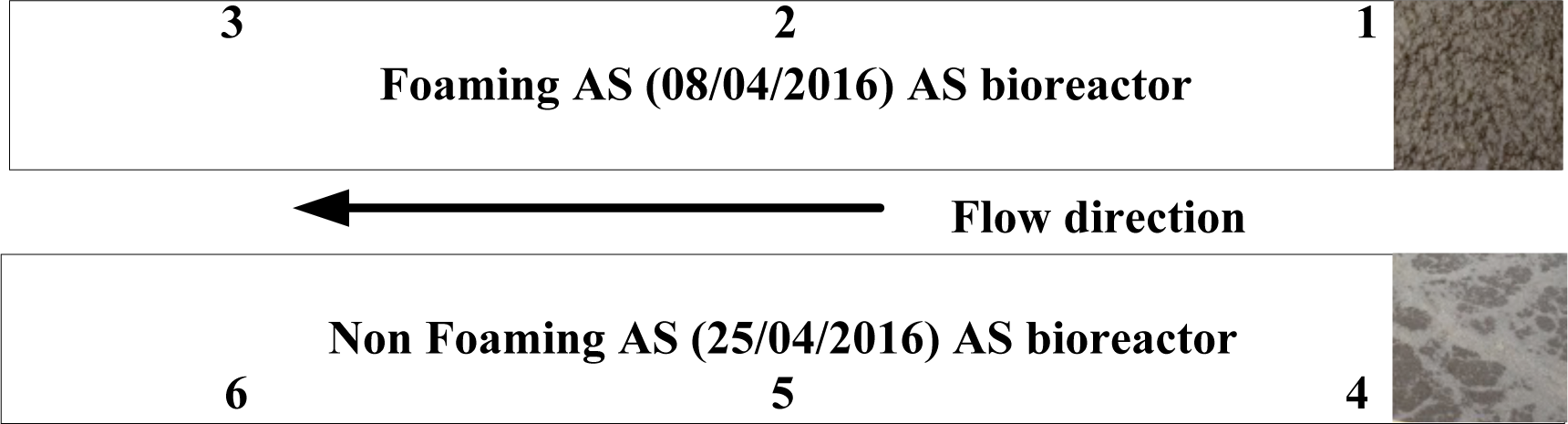
Samples were collected for foaming activated sludge at 08/04/2016 and non-foaming activated sludge at 25/04/2016 alongside the bioreactor at Shatin wastewater treatment plant.

### Whole DNA, total RNA extraction, removal of ribosomal RNA, cDNA synthesis and next generation sequencing

FAST DNA Kit was used to extract total DNA from 1 mL mixed AS samples. RNeasy Mini was used to extract the total RNA from 0.5 mL AS stored in liquid nitrogen. The extracted RNA was then processed by DNase I to eliminate the DNA in the RNA samples. Then both Illumina Ribo-Zero rRNA removal KIT (Bacteria) and Ribo-Zero rRNA removal KIT (Human/Mouse/Rat) was applied for each sample to remove rRNA from Prokaryote and Eukaryote respectively in order to get the total clean non-ribosomal RNA. Generally, metatranscriptome rRNA depletion was only used the Ribo-Zero for Bacteria, in this study, the addition of Eukaryote rRNA removal was due to a fact that by only using the Ribo-Zero Bacteria rRNA removal Kit for AS, there was still over half of RNA were rRNA from Eukaryote (our previous experiment, data unpublished). To get more non-rRNA, the Ribo-Zero rRNA Kit to remove Eukaryote was also used. RNA then was fragmented into 170 bps library and was reverse-transcribed to construct cDNA library for sequencing. The quality of DNA and RNA were assessed with Agilent 2100 Bioanalyzer (Agilent Technologies, Palo Alto, CA, USA). All the samples was sent to sequence, considering the complexity of AS and the aims of this study to detect the expression of low abundance gene, we gave each sample a very deep sequencing depth which doubled the sequencing depth in previous studies. All the samples were sequenced with Hiseq 4000 in BGI-ShenZhen. DNA samples with PE-150 with library size of 300 bps. And RNA with PE101 of library size 170 bps.

### Bioinformatics analysis

Quality filtering was firstly performed on DNA and RNA reads to keep only high quality reads using trimmomatic v1.04 (Bolger et al. 2014). DNA datasets were pooled together and assembled by CLC Genomics Workbench 6.5.3 (CLC Bio, Aarhus, Denmark, https://www.qiagenbioinformatics.com/) with default parameters. Finally, 1,430,611 contigs with length over 100 bps (N50, 2,416 bps; 2,457,704,443 bps length in total) were obtained and 74.5% of reads could be mapped back to these Contigs. All these contigs were sent to predict genes with Prodigal (version 1.5) (Hyatt et al. 2010) using `-meta` parameter and finally 3,234,330 genes were obtained. By removing exactly the same genes using USEARCH (version 8.0.1623) (Edgar 2010) unique command (parameters ‐fastx_uniques), 3,234,246 million genes were kept; this set was defined as ‘unique gene set’. Reads were mapped back to the contig set and ‘unique gene set’ to obtain reads coverage matrixes for contigs and genes. The matrix of genes was finally normalized to cell numbers. For metatranscriptome samples, after quality filtering, the SortMeRNAv1.9 was used to remove all the possible ribosomal RNA by aligning to six databases of bacteria, archaea and eukaryotic small and large subunits (Kopylova et al. 2012). RNA reads for each sample were then mapped back to the `unique gene set` to get the transcript coverage for each gene with CLC genomic workbench 6.5.3 using parameters of gap penalty 2, gap extension 3, length fraction 0.8 and similarity at least 0.9.

Taxonomy composition of the metagenome was generated with MEGAN6 (Huson et al. 2015). In detail, all genes were aligned to NCBI NR database (version 201603) with diamondv1.09 (Buchfink et al. 2015) to find out the homology proteins. To each gene, the local common ancestors (LCA) were applied using the taxonomy information of the hit NR protein in NCBI taxonomy database (Acland et al. 2014) and then this gene was annotated with the common ancestor taxonomy. We further processed the NCBI taxonomy annotation results to remove those subdivisions and subgroups to format the annotation to 7 levels from kingdom to species. Among total 3,234,246 unique genes predicted, 2,348,907 could be aligned to NR database. The remaining 885,339 (27.3%) genes could not be annotated with the NR database. The abundance of each taxon was a sum of all the annotated genes under that taxon in every sample. Antibiotic resistant genes (ARGs) were annotated with SARG database which contained a type-subtype structure annotation (Yang et al. 2016). Metal resistance genes (MRGs) were detected by aligning the “unique gene set” to the MRG database (Li et al. 2017). Absolute abundance and transcript was determined by AQMM.

## Declarations

### Data availability

The metagenome and metatranscriptome raw data were deposited in NCBI SRA under accession number XXX.

### Analyzing document

The analyzing document for the whole data analysis and simulation process could be accessed from https://github.com/biofuture/aqmm/blob/master/Analysing_document.txt

## Acknowledgements

The authors would like to thank GRF of Hong Kong for financial support (172099/14E). Xiaotao Jiang would like to thank The University of Hong Kong for the postgraduate scholarship, and Dr. Ke Yu, Dr. Li-Guan, and Dr. Andong, Li, would like to thank The University of Hong Kong for postdoc fellowship.

## Contributions

T. Zhang and X.-T. Jiang design the study of quantification. X.-T. Jiang developed the software and performed the wet-lab and simulation experiments. X.-T. Jiang performed the bioinformatics analyses. X.-T. Jiang, A.D. Li and K. Y. did the DNA and RNA extraction experiment. L.-G. Li did the MRG analyses. T. Zhang and X.-T. Jiang wrote the manuscript. T. Zhang, X.-T. Jiang, A.D. Li, L.G. Li and X.L. Yin revised the manuscript.

## Reference

Acland, A., Agarwala, R., Barrett, T., Beck, J., Benson, D.A., Bollin, C., Bolton, E., Bryant, S.H., Canese, K. and Church, D.M. (2014) Database resources of the national center for biotechnology information. Nucleic acids research 42(Database issue), D7.

Alexander, H., Jenkins, B.D., Rynearson, T.A. and Dyhrman, S.T. (2015) Metatranscriptome analyses indicate resource partitioning between diatoms in the field. Proceedings of the National Academy of Sciences 112(17), E2182–E2190.

Becker-André, M. and Hahlbrock, K. (1989) Absolute mRNA quantification using the polymerase chain reaction (PCR). A novel approach by a PCR aided transcipt titration assay (PATTY). Nucleic acids research 17(22), 9437–9446.

Bolger, A.M., Lohse, M. and Usadel, B. (2014) Trimmomatic: a flexible trimmer for Illumina sequence data. Bioinformatics, btu170.

Buchfink, B., Xie, C. and Huson, D.H. (2015) Fast and sensitive protein alignment using DIAMOND. Nature Methods 12(1), 59–60.

Conesa, A., Madrigal, P., Tarazona, S., Gomez-Cabrero, D., Cervera, A., McPherson, A., Szczesniak, M.W., Gaffney, D.J., Elo, L.L. and Zhang, X. (2016) A survey of best practices for RNA-seq data analysis. Genome biology 17(1), 13.

Edgar, R.C. (2010) Search and clustering orders of magnitude faster than BLAST. Bioinformatics 26(19), 2460–2461.

Foladori, P., Bruni, L., Tamburini, S. and Ziglio, G. (2010) Direct quantification of bacterial biomass in influent, effluent and activated sludge of wastewater treatment plants by using flow cytometry. Water Research 44(13), 3807–3818.

Gifford, S.M., Sharma, S., Rinta-Kanto, J.M. and Moran, M.A. (2011) Quantitative analysis of a deeply sequenced marine microbial metatranscriptome. The ISME journal 5(3), 461–472.

Huson, D., Beier, S., Buchfink, B., Flade, I., Górska, A., El-Hadidi, M., Mitra, S., Ruscheweyh, H.-J. and Tappu, R. (2015) MEGAN6-Microbiome analysis involving hundreds of samples and billions of reads, preparation.

Hyatt, D., Chen, G.L., Locascio, P.F., Land, M.L., Larimer, F.W. and Hauser, L.J. (2010) Prodigal: prokaryotic gene recognition and translation initiation site identification. BMC bioinformatics 11(1), 119.

Jorth, P., Turner, K.H., Gumus, P., Nizam, N., Buduneli, N. and Whiteley, M. (2014) Metatranscriptomics of the human oral microbiome during health and disease. MBio 5(2), e01012–o01014.

Kanno, J., Aisaki, K.-i., Igarashi, K., Nakatsu, N., Ono, A., Kodama, Y. and Nagao, T. (2006) “ Per cell” normalization method for mRNA measurement by quantitative PCR and microarrays. BMC genomics 7(1), 64.

Klingenberg, H. and Meinicke, P. (2017) How To Normalize Metatranscriptomic Count Data For Differential Expression Analysis. bioRxiv, 134650.

Kopylova, E., Noé, L. and Touzet, H. (2012) SortMeRNA: fast and accurate filtering of ribosomal RNAs in metatranscriptomic data. Bioinformatics 28(24), 3211–3217.

Li, L.G., Xia, Y. and Zhang, T. (2017) Co-occurrence of antibiotic and metal resistance genes revealed in complete genome collection. Isme Journal 11(3), 651–662.

Love, M.I., Huber, W. and Anders, S. (2014) Moderated estimation of fold change and dispersion for RNA-seq data with DESeq2. Genome biology 15(12), 550.

Manti, A., Boi, P., Falcioni, T., Canonico, B., Ventura, A., Sisti, D., Pianetti, A., Balsamo, M. and Papa, S. (2008) Bacterial cell monitoring in wastewater treatment plants by flow cytometry. Water Environment Research 80(4), 346–354.

Martinez, X., Pozuelo, M., Pascal, V., Campos, D., Gut, I., Gut, M., Azpiroz, F., Guarner, F. and Manichanh, C. (2016) MetaTrans: an open-source pipeline for metatranscriptomics. Scientific reports 6, 26447.

Mason, O.U., Hazen, T.C., Borglin, S., Chain, P.S., Dubinsky, E.A., Fortney, J.L., Han, J., Holman, H.-Y.N., Hultman, J. and Lamendella, R. (2012) Metagenome, metatranscriptome and single-cell sequencing reveal microbial response to Deepwater Horizon oil spill. The ISME journal 6(9), 1715–1727.

Moran, M.A., Satinsky, B., Gifford, S.M., Luo, H., Rivers, A., Chan, L.-K., Meng, J., Durham, B.P., Shen, C. and Varaljay, V.A. (2013) Sizing up metatranscriptomics. The ISME journal 7(2), 237.

Mortazavi, A., Williams, B.A., McCue, K., Schaeffer, L. and Wold, B. (2008) Mapping and quantifying mammalian transcriptomes by RNA-Seq. Nature methods 5(7), 621–628.

Mumy, K.L. and Findlay, R.H. (2004) Convenient determination of DNA extraction efficiency using an external DNA recovery standard and quantitative-competitive PCR. Journal of Microbiological Methods 57(2), 259–268.

Narayanasamy, S., Jarosz, Y., Muller, E.E., Heintz-Buschart, A., Herold, M., Kaysen, A., Laczny, C.C., Pinel, N., May, P. and Wilmes, P. (2016) IMP: a pipeline for reproducible reference-independent integrated metagenomic and metatranscriptomic analyses. Genome biology 17(1), 260.

Nayfach, S. and Pollard, K.S. (2015) Average genome size estimation improves comparative metagenomics and sheds light on the functional ecology of the human microbiome. Genome biology 16(1), 51.

Ni, Y., Li, J. and Panagiotou, G. (2016) COMAN: a web server for comprehensive metatranscriptomics analysis. BMC genomics 17(1), 622.

Oyserman, B.O., Noguera, D.R., del Rio, T.G., Tringe, S.G. and McMahon, K.D. (2015) Metatranscriptomic insights on gene expression and regulatory controls in Candidatus Accumulibacter phosphatis. The ISME journal.

Rapaport, F., Khanin, R., Liang, Y., Pirun, M., Krek, A., Zumbo, P., Mason, C.E., Socci, N.D. and Betel, D. (2013) Comprehensive evaluation of differential gene expression analysis methods for RNA-seq data. Genome biology 14(9), 3158.

Robinson, M.D., McCarthy, D.J. and Smyth, G.K. (2010) edgeR: a Bioconductor package for differential expression analysis of digital gene expression data. Bioinformatics 26(1), 139–140.

Robinson, M.D. and Oshlack, A. (2010) A scaling normalization method for differential expression analysis of RNA-seq data. Genome biology 11(3), R25.

Shi, Y.M., Tyson, G.W. and DeLong, E.F. (2009) Metatranscriptomics reveals unique microbial small RNAs in the ocean’s water column. nature 459(7244), 266–U154.

Stark, L., Giersch, T. and Wünschiers, R. (2014) Efficiency of RNA extraction from selected bacteria in the context of biogas production and metatranscriptomics. Anaerobe 29, 85–90.

Turner, T.R., Ramakrishnan, K., Walshaw, J., Heavens, D., Alston, M., Swarbreck, D., Osbourn, A., Grant, A. and Poole, P.S. (2013) Comparative metatranscriptomics reveals kingdom level changes in the rhizosphere microbiome of plants. The ISME journal 7(12), 2248–2258.

Wagner, G.P., Kin, K. and Lynch, V.J. (2012) Measurement of mRNA abundance using RNA-seq data: RPKM measure is inconsistent among samples. Theory in Biosciences 131(4), 281–285.

Westreich, S.T., Korf, I., Mills, D.A. and Lemay, D.G. (2016) SAMSA: a comprehensive metatranscriptome analysis pipeline. BMC bioinformatics 17(1), 399.

Whelan, J.A., Russell, N.B. and Whelan, M.A. (2003) A method for the absolute quantification of cDNA using real-time PCR. Journal of immunological methods 278(1), 261–269.

Yang, Y., Jiang, X.T., Chai, B.L., Ma, L.P., Li, B., Zhang, A.N., Cole, J.R., Tiedje, J.M. and Zhang, T. (2016) ARGs-OAP: online analysis pipeline for antibiotic resistance genes detection from metagenomic data using an integrated structured ARG-database. Bioinformatics 32(15), 2346–2351.

Yu, K. and Zhang, T. (2012) Metagenomic and metatranscriptomic analysis of microbial community structure and gene expression of activated sludge. Plos One 7(5), e38183.

